# Single-cell entropy for accurate estimation of differentiation potency from a cell’s transcriptome

**DOI:** 10.1101/084202

**Authors:** Andrew E Teschendorff

**Affiliations:** CAS Key Laboratory of Computational Biology, CAS-MPG Partner Institute for Computational Biology, 320 Yue Yang Road, Shanghai 200031, China.; Department of Women’s Cancer, University College London, 74 Huntley Street, London WC1E 6AU, United Kingdom.; Statistical Cancer Genomics, Paul O’Gorman Building, UCL Cancer Institute, University College London, 72 Huntley Street, London WC1E 6BT, United Kingdom.

**Keywords:** Single-Cell, RNA-Seq, Stem-Cell, Differentiation, Cancer, Entropy

## Abstract

The ability to quantify differentiation potential of single cells is a task of critical importance for single-cell studies. So far however, there is no robust general molecular correlate of differentiation potential at the single cell level. Here we show that differentiation potency of a single cell can be approximated by computing the signaling promiscuity, or entropy, of a cell’s transcriptomic profile in the context of a cellular interaction network, without the need for model training or feature selection. We validate signaling entropy in over 7,000 single cell RNA-Seq profiles, representing all main differentiation stages, including time-course data. We develop a novel algorithm called <u>S</u>ingle <u>C</u>ell <u>Ent</u>ropy (SCENT), which correctly identifies known cell subpopulations of varying potency, enabling reconstruction of cell-lineage trajectories. By comparing bulk to single cell data, SCENT reveals that expression heterogeneity within single cell populations is regulated, pointing towards the importance of cell-cell interactions. In the context of cancer, SCENT can identify drug resistant cancer stem-cell phenotypes, including those obtained from circulating tumor cells. In summary, SCENT can directly estimate the differentiation potency and plasticity of single-cells, allowing unbiased quantification of intercellular heterogeneity, and providing a means to identify normal and cancer stem cell phenotypes.

**Software Availability:** SCENT is freely available as an R-package from github: https://github.com/aet21/SCENT

---

One of the most important tasks in single-cell RNA-sequencing studies is the identification and quantification of intercellular transcriptomic heterogeneity [1–4]. Although some of the observed heterogeneity represents stochastic noise, a substantial component of intercellular variation has been shown to be of functional importance [1, 5–8]. Very often, this biologically relevant heterogeneity can be attributed to cells occupying states of different potency or plasticity. Thus, quantification of differentiation potency, or more generally functional plasticity, at the single-cell level is of paramount importance. However, currently there is no concrete theoretical and computational model for estimating such plasticity at the single cell level.

Here we make significant progress towards addressing this challenge. We propose a very general model for estimating cellular plasticity. A key feature of this model is the computation of signaling entropy [9], which quantifies the degree of uncertainty, or promiscuity, of a cell’s gene expression levels in the context of a cellular interaction network. We show that signaling entropy provides an excellent and robust proxy to the differentiation potential of a cell in Waddington’s epigenetic landscape [10], and further provides a framework in which to understand the overall differentiation potency and transcriptomic heterogeneity of a cell population in terms of single-cell potencies. Attesting to its general nature and broad applicability, we compute and validate signaling entropy in over 7000 single cells of variable degrees of differentiation potency and phenotypic plasticity, including time-course differentiation data, neoplastic cells and circulating tumor cells (CTCs). This extends entropy concepts that we have previously demonstrated to work on bulk tissue data [9, 11–13] to the single-cell level. Based on signaling entropy, we develop a novel algorithm called SCENT (<u>S</u>ingle <u>C</u>ell <u>Ent</u>ropy), which can be used to identify and quantify biologically relevant expression heterogeneity in single-cell populations, as well as to reconstruct cell-lineage trajectories from time-course data. In this regard, SCENT differs substantially from other single-cell algorithms like Monocle [14], MPath [15], SCUBA [16], Diffusion Pseudotime [17] or StemID [18], in that it uses single-cell entropy to independently order single cells in pseudo-time (i.e. differentiation potency), without the need for feature selection or clustering.

## Results

### Single-cell entropy as a proxy to the differentiation potential of single cells in Waddington’s landscape

A pluripotent cell (by definition endowed with the capacity to differentiate into effectively all major cell-lineages) does not express a preference for any particular lineage, thus requiring a similar basal activity of all lineage-specifying transcription factors [9, 19]. Viewing a cell’s choice to commit to a particular lineage as a probabilistic process, pluripotency can therefore be characterized by a state of high uncertainty, or entropy, because all lineage-choices are equally likely (**Fig.1A**). In contrast, for a differentiated cell, or for a cell committed to a particular lineage, signaling uncertainty/entropy is reduced, as this requires activation of a specific signaling pathway reflecting that lineage choice (**Fig.1A)**. Thus, a measure of global signaling entropy, if computable, could provide us with a relatively good proxy of a cell’s overall differentiation potential. Here we propose that signaling entropy can be estimated *in-silico* by integrating a cell’s transcriptomic profile with a high quality protein-protein-interaction (PPI) network to define a cell-specific stochastic “random-walk” matrix from which a global signaling entropy (abbreviated as SR) can then be computed (**Fig.1A-B, Online Methods**). It can be shown that signaling entropy is, in effect, the correlation of a cell’s transcriptomic profile with the connectivity profile of the corresponding proteins in the PPI network (**Online Methods**). Underlying our model is therefore the assumption that highly connected proteins are more likely to be highly expressed under conditions in which the requirement for a cell’s phenotypic plasticity is high (e.g. as in a pluripotent state).

**Figure-1.**
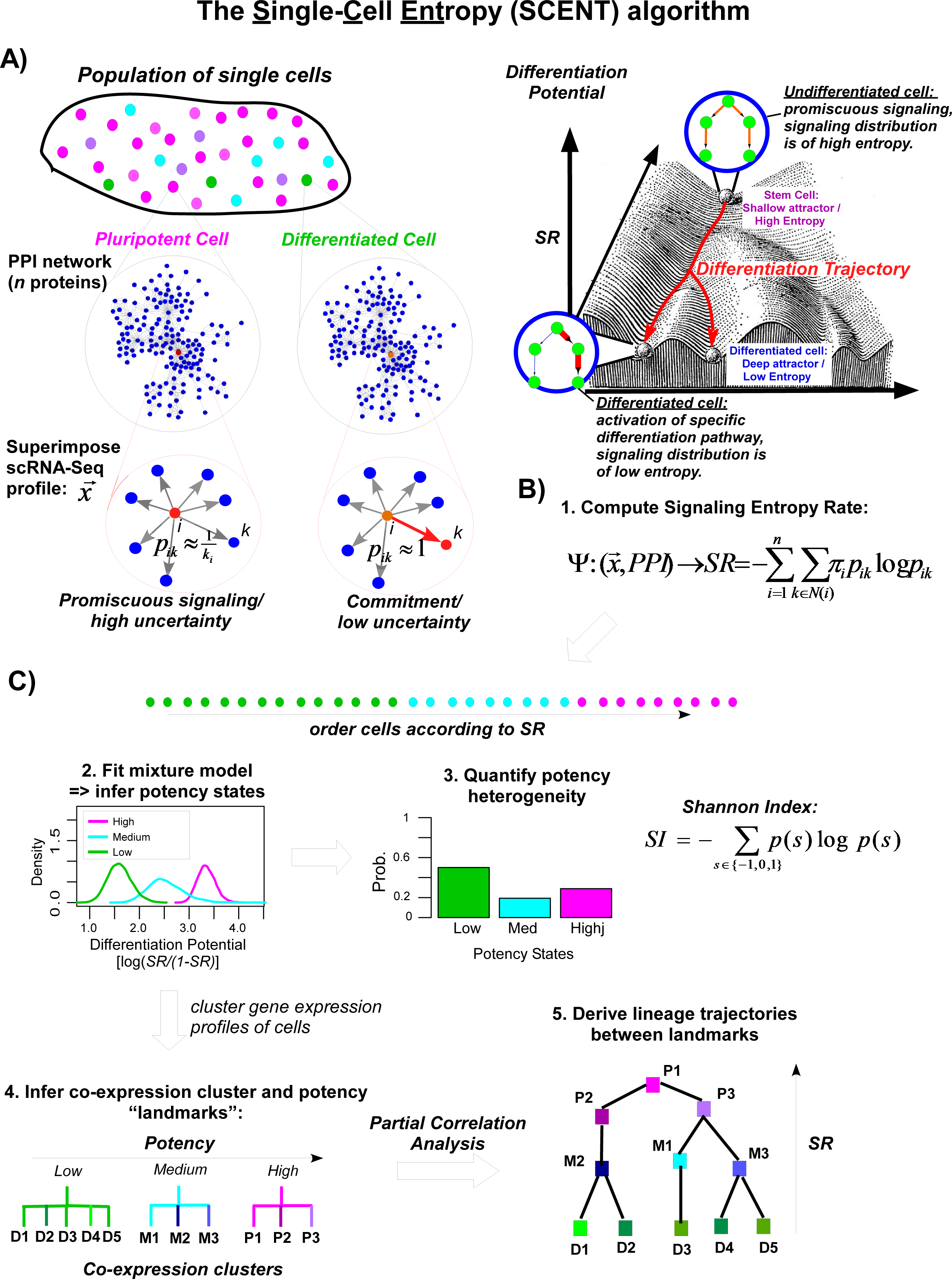
The Single-Cell Entropy (SCENT) algorithm. A) Signaling entropy of single cells as a proxy to their differentiation potential in Waddington’s landscape. Depicted on the left is a population of cells with cells occupying either a pluripotent (magenta), a progenitor (cyan) or a differentiated state (green). The potency state of each cell is determined by a complex function of the transcriptomic profile of the cell. In a pluripotent state, there is high demand for phenotypic plasticity, and so promiscuous signaling proteins (i.e those of high connectivity) are highly expressed (red colored node) with all major differentiation pathways kept at a similar basal activity level (grey edges). The probability of signaling between protein *i* and *k, p*_*ik*_, is therefore 1/*k*_*i*_ where *k*_*i*_ is the connectivity of protein *i* in the network. In a differentiated state, commitment to a specific lineage (activation of a specific signaling pathway shown by red colored node) means that most *p*_*ij*_~0, except when *j=k*, so that *p*_*ik*_~1. On the right we depict a cartoon of Waddington’s epigenetic landscape, illustrating the same concept. **B) Estimation of signaling entropy.** Approximation of the differentiation potential of a single cell by computation of the signaling entropy rate (SR) over all the genes/proteins in the network, where *π* is the invariant measure (steady-state probability). **C) Quantification of intercellular heterogeneity and reconstruction of lineage trajectories.** Estimation of signaling entropy at the single-cell level across a population of cells, allows the distribution of potency states in the population to be determined through Bayes mixture modelling which infers the optimal number of potency states. From this, the heterogeneity of potency states in a cell population is computed using Shannon’s Index. To infer lineage trajectories, SCENT uses a clustering algorithm over dimensionally reduced scRNA-Seq profiles to infer co-expression clusters of cells. Dual assignment of cells to a potency state and co-expression cluster allows the identification of landmarks as bi-clusters in potency-coexpression space. Finally, partial correlations between the expression profiles of the landmarks are used to infer a lineage trajectory network diagram linking cell clusters according to expression similarity, with their height or elevation determined by their potency (signaling entropy).

### Validation of single-cell entropy as a measure of differentiation potency

To test that signaling entropy correlates with differentiation potency, we first estimated it for 1018 single-cell RNA-seq profiles generated by Chu et al [20], which included pluripotent human embryonic stem cells (hESCs) and hESC-derived progenitor cells representing the 3 main germ-layers (endoderm, mesoderm and ectoderm) (“Chu et al set”, **SI table S1, Online Methods**). In detail, these were 374 cells from two hESC lines (H1 & H9), 173 neural progenitor cells (NPCs), 138 definite endoderm progenitors (DEPs), 105 endothelial cells representing mesoderm derivatives, as well as 69 trophoblast (TB) cells and 148 human foreskin fibroblasts (HFFs). Confirming our hypothesis, pluripotent hESCs attained the highest signaling entropy values, followed by multipotent cells (NPCs, DEPs), and with less multipotent HFFs, TBs and ECs attaining the lowest values (**Fig.2A**). Differences were highly statistically significant, with DEPs exhibiting significantly lower entropy values than hESCs (Wilcoxon rank sum P<1e-50 (**Fig.2A**). Likewise, TBs exhibited lower entropy than hESCs (P<1e-50), but higher than HFFs (P<1e-7) (**Fig.2A**). Importantly, signaling entropy correlated very strongly with a pluripotency score obtained using a previously published pluripotency gene expression signature [21] (Spearman Correlation = 0.91, P<1e-500, **Fig.2B**, **Online Methods**). In all, signaling entropy provided a highly accurate discriminator of pluripotency versus non-pluripotency at the single cell level (AUC=0.96, Wilcoxon test P<1e-300, **Fig.2C**). We note that in contrast with pluripotency expression signatures, this strong association with pluripotency was obtained *without the need for any feature selection or training*.

**Figure-2.**
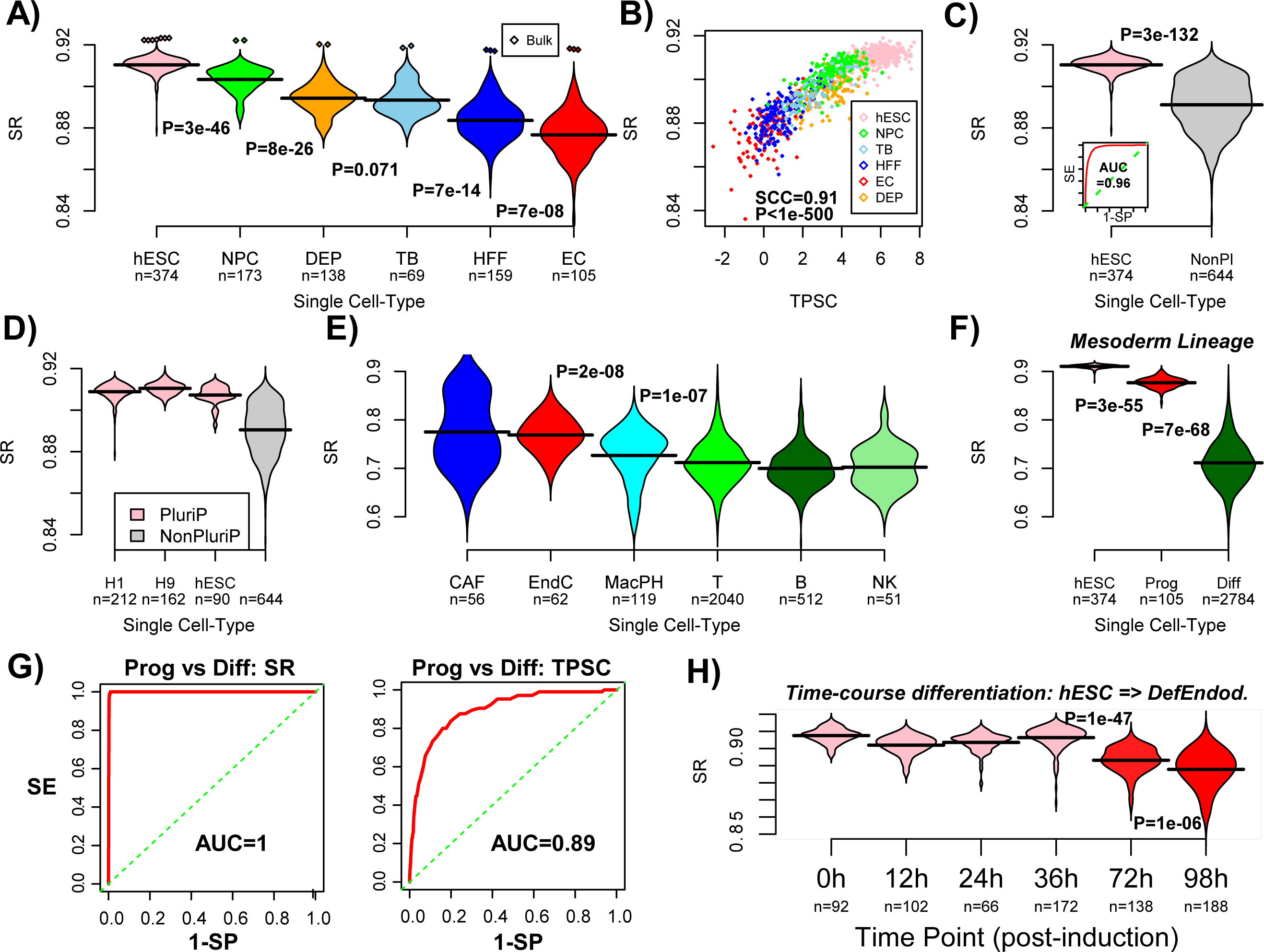
Signaling entropy correlates with differentiation potency of single cells. A) Violin plots of the signaling entropy (SR) against cell-type (hESC=human embryonic stem cells, NPC=neural progenitor cells, DEP=definite endoderm progenitors, TB=trophoblast cells, HFF=human foreskin fibroblasts, EC=endothelial cells (mesoderm progenitor derivatives)). Number of single cells in each class is indicated. Total number is 1018. Wilcoxon rank sum test P-values between each cell-type (ranked in decreasing order of SR) are given. Diamond shaped data points correspond to the matched bulk samples. **B)** Scatterplot of the signaling entropy (SR, y-axis) against an independent mRNA expression based pluripotency score (TPSC, x-axis) for all 1018 single cells. Cell-type is indicated by color. Spearman Correlation Coefficient (SCC) and associated P-value are given. **C)** Violin plot comparing the signaling entropy (SR) between the hESCs and all other (non-pluripotent) cells. P-value is from a Wilcoxon rank sum test. Inlet figure is the associated ROC curve, which includes the AUC value. **D)** As C), but now splitting the hESCs into cells from H1 and H9 lines, and including an additional independent set of 90 single hESCs profiled with a different NGS platform. **E)** Violin plot of signaling entropy (SR) values for non-malignant single cells found in the microenvironment of melanomas. Number of single cells of each cell-type are given (CAF=cancer associated fibroblasts, EndC=endothelial cells, MacPH=macrophages, T=T-cells, B=B-cells, NK=natural killer cells). Wilcoxon rank sum test P-values between EndC and MacPH, and between MacPH and all lymphocytes are given. **F)** Signaling entropy (SR) as a function of differentiation stage within the mesoderm lineage. Differentiation stages include hESCs (pluripotent), mesoderm progenitors of endothelial cells (multipotent) and differentiated endothelial and white blood cells. Wilcoxon rank sum test P-values between successive stages are given. **G)** ROC curves and AUC values for discriminating the progenitor and differentiated cells within the mesoderm lineage for signaling entropy (SR) and the t-test pluripotency score (TPSC). **H)** Signaling entropy (SR, y-axis) as a function of time in a single-cell time course differentiation experiment, starting from hESCs at time=0h (time of differentiation induction) into definite endoderm (which occurs from 72h onwards). Number of single cells measured at each time point is given. Wilcoxon rank sum test P-values between the first 4 time points and 72h, and between 72h and 98h are given.

To further test the general validity and robustness of signaling entropy we computed it for scRNA-Seq profiles of 3256 non-malignant cells derived from the microenvironment of 19 melanomas (Melanoma set, [22], **SI table S1**). Cells profiled included T-cells, B-cells, natural-killer (NK) cells, macrophages, fully differentiated endothelial cells and cancer-associated fibroblasts (CAFs). For a given cell-type and individual, variation between single cells was substantial and similar to the variation seen between individuals (**SI fig.S1**). Mean entropy values however, were generally stable, showing little inter-individual variation, except for T-cells from 4 out of 15 patients, which exhibited a distinctively different distribution (**SI fig.S1**). In order to assess overall trends, we pooled the single-cell entropy data from all patients together, which confirmed that all lymphocytes (T-cells, B-cells and NK-cells) had similar average signaling entropy values (**Fig.2E**). Intra-tumor macrophages, which are derived from monocytes, exhibited a marginally higher signaling entropy (**Fig.2E**). The highest signaling entropy values were attained by endothelial cells and CAFs (**Fig.2E**), consistent with their known high phenotypic plasticity [23–26]. Importantly, the entropy values for all of these non-malignant differentiated cell-types were distinctively lower compared to those of hESCs and progenitor cells from Chu et al (**Figs.2A & 2E**), consistent with the fact that hESCs and progenitors have much higher differentiation potency. To test this formally, we compared hESCs, mesoderm progenitors, and terminally differentiated cells within the mesoderm lineage (which included all endothelial cells and lymphocytes), which revealed a consistent decrease in signaling entropy between all three potency states (Wilcoxon rank test P<1e-50, **Fig.2F**). Of note, signaling entropy could discriminate progenitor and differentiated cells better than the score derived from the pluripotency gene expression signature [21], attesting to its increased robustness as a general measure of differentiation potency (**Fig.2G**, **SI fig.S2**).

Next, we assessed signaling entropy in the context of a time-course differentiation experiment, whereby hESCs were induced to differentiate into definite endoderm progenitors via the mesoendoderm intermediate [27]. scRNA-Seq for a total of 758 single cells, obtained at 6 timepoints, including origin, 12, 24, 36, 72 and 96 hours post-induction were available (**Online Methods**) [27]. We observed that single cell entropies exhibited a particular large decrease only after 72 hours (**Fig.2H**), consistent with previous knowledge that differentiation into definite endoderm occurs around 3-4 days after induction [27].

### Signaling entropy is robust to choice of PPI network and NGS platform

We verified that signaling entropy is robust to the choice of PPI network (**SI fig.S3**). This robustness to the network stems from the fact that signaling entropy depends mainly on the relative connectivity of the proteins in the network (**SI fig.S4A**). Importantly, signaling entropy lost its power to discriminate pluripotent from non-pluripotent cells if expression values were randomly reshuffled over the network (**SI fig.S4B-C**), demonstrating that features such as pluripotency are encoded in a subtle positive correlation between expression levels and connectivity. In order to test the robustness of signaling entropy across independent studies, we analyzed scRNA-Seq data for an independent set of single cell hESCs derived from the primary outgrowth of the inner cell mass (“hESC set” [28], **SI table S1**). Obtained signaling entropy values were most similar to those of single cells derived from the H1 and H9 hESC lines, confirming the robustness of signaling entropy across different studies and next-generation sequencing platforms (**Fig.2D, SI table S1**).

### Non-linear association between single cell entropy and cell-cycle phase

A major source of variation in scRNA-Seq data is cell-cycle phase [22, 29]. We explored the relation between signaling entropy and cell-cycle phase in a large scRNA-Seq dataset encompassing 3256 non-malignant and 1257 cancer cells derived from the microenvironment of melanomas (Melanoma set, [22], **SI table S1**). A cycling score for both G1-S and G2-M phases and for each cell was obtained using a validated procedure [22, 29, 30] and compared to signaling entropy, which revealed a strong yet highly non-linear correlation (**SI fig.S5**). Specifically, we observed that cells with a low signaling entropy were never found in either the G1-S or G2-M phase (**SI fig.S5**). In contrast, cells with high signaling entropy could be found in either a cycling or non-cycling phase. These results are consistent with the view that cycling-cells must increase expression of promiscuous signaling proteins and hence exhibit an increased signaling entropy.

### Quantification of inter-cellular expression heterogeneity with SCENT

Given that signaling entropy correlates with differentiation potency, we used it to develop the SCENT algorithm (**Fig.1C**). Briefly, the SCENT algorithm uses the estimated signaling entropies of single cells to derive the distribution of discrete potency states across the cell population (**Fig.1C, Online Methods**). Thus, SCENT can be used to quantify expression heterogeneity at the level of potency. In addition, SCENT can be used to directly order single cells in pseudo-time [14] to facilitate reconstruction of lineage trajectories. A key feature of SCENT is the assignment of each cell to a unique potency state and co-expression cluster, which results in the identification of potency-clusters (which we call “landmarks”), through which lineage trajectories are then inferred (**Online Methods**).

To test SCENT, we applied it to the scRNA-Seq data from Chu et al, a non-time course single-cell experiment, which includes hESCs and progenitor cell populations (**SI table S1**). SCENT correctly predicted a parsimonious two potency state model, with a high potency pluripotent state and a lower potency non-pluripotent progenitor-like state (**Fig.3A**). Interestingly, a small fraction (approximately 4%) of the single hESCs were deemed to be non-pluripotent cells (**Fig.3B**), consistent with previous observations that pluripotent cell populations contain cells that are already primed for differentiation into specific lineages [5, 6]. Supporting this further, these non-pluripotent “hESC” cells exhibited lower cycling-scores and higher expression levels of neural (*HES1/SOX2*) and mesoderm (*PECAM1*) stem-cell markers, compared to the pluripotent hESC cells (**SI fig.S6**). Whereas all HFFs and ECs were deemed non-pluripotent, definite endoderm progenitors (DEPs), TBs and NPCs exhibited mixed proportions, with NPCs exhibiting approximately equal numbers of pluripotent and non-pluripotent cells (**Fig.3B**). Correspondingly, the Shannon index, which quantifies the level of heterogeneity in potency, was highest for the NPC population (**Fig.3C**). In total, SCENT predicted 6 co-expression clusters, which combined with the two potency states, resulted in a total of 7 landmark clusters (**Fig.3D**). These landmarks correlated very strongly with cell-type, with only NPCs being distributed across two landmarks of different potency (**Fig.3E**). SCENT correctly inferred a lineage trajectory between the high potency NPC subpopulation and its lower potency counterpart, as well as a trajectory between hESCs and DEPs (**Fig.3F**). The other cell-types exhibited lower entropies (**Fig.2B & Fig.3F**), and correspondingly did not exhibit a direct trajectory to hESCs, suggesting several intermediate states which were not sampled in this experiment.

**Figure-3.**
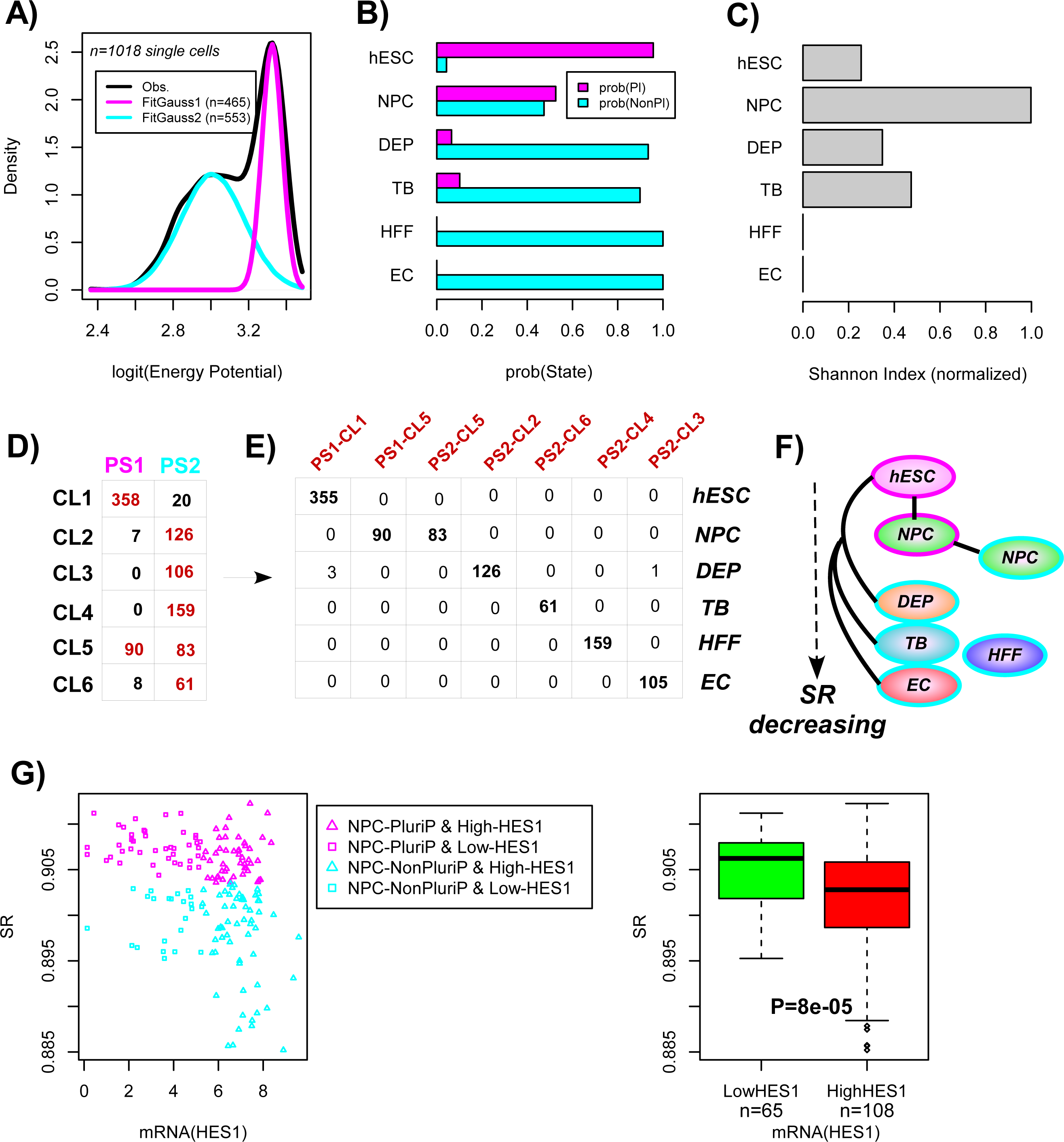
SCENT identifies single cell subpopulations of biological significance. A) Fitted Gaussian mixture model to the signaling entropies of 1018 single cells (scRNA-Seq data from Chu et al) using a logit scale for the signaling entropies (x-axis, log2[SR/(1-SR)]). BIC predicted only 2-states: a high energy/entropy pluripotent state (magenta-PS1) and a lower-energy non-pluripotent state (cyan-PS2). Number of cells categorized into each state is indicated in plot. **B)** Barplot comparing, for each cell-type, the probability that a cell from this cell population is in the pluripotent (prob(Pl)) or non-pluripotent state (probe(NonPl). Cell-types include human embryonic stem cells (hESCs), neural progenitor cells (NPCs), definite endoderm progenitors (DEPs), trophoblast cells (TBs), human foreskin fibroblasts (HFFs) and endothelial cells (ECs). **C)** Barplot of the corresponding Shannon Index for each cell-population type. **D)** Distribution of single cell numbers between inferred potency states and co-expression clusters, as predicted by SCENT. In brown, we indicate “landmark clusters” which contain at least 5% of the total number of single cells. **E)** Distribution of single cell-types among the 7 landmark clusters. **F)** Inferred lineage trajectories between the 7 landmarks which map to cell-types. Border color indicates potency state: magenta=PS1, cyan=PS2. **G) Left panel:** Scatterplot of signaling entropy (SR) vs mRNA expression level of a neural stem/progenitor cell marker, HES1, for all NPCs. NPCs categorized as pluripotent are shown in magenta, NPCs categorized into a non-pluripotent state are shown in cyan. NPCs of high and low HES1 expression (as inferred using a partition-around-medoids algorithm with k=2) are indicated with triangles and squares, respectively. **Right panel:** Corresponding boxplot comparing the differentiation potency (SR) of NPCs with low vs. high HES1 expression. P-value is from a one-tailed Wilcoxon rank sum test.

To ascertain the biological significance of the two NPC subpopulations (**Fig.3B,E,F**), we first verified that the NPCs deemed pluripotent did indeed have a higher pluripotency score (**SI Fig.7A**), as assessed using the independent pluripotency gene expression signature from Palmer et al [21]. We further reasoned that well-known transcription factors marking neural stem/progenitor cells, such as HES1, would be expressed at a much lower level in the NPCs deemed pluripotent compared to the non-pluripotent ones, since the latter are more likely to represent *bona-fide* NPCs. Confirming this, NPCs with low HES1 expression exhibited higher differentiation potential than NPCs with high HES1 expression (Wilcoxon rank sum test P<0.0001, **Fig.3G**). Similar results were evident for other neural progenitor/stem cell markers such as PAX6 and SOX2 (**SI fig.S7B**). Of note, NPCs expressing the lowest levels of PAX6, HES1 or SOX2 were generally always classified by SCENT into a pluripotent-like state (**Fig.3G, SI fig.S7B**). Thus, these results indicate that SCENT provides a biologically meaningful characterization of intercellular transcriptomic heterogeneity.

### SCENT reconstructs lineage trajectories of human myoblast differentiation

We next tested SCENT in the context of a differentiation experiment of human myoblasts [14], involving skeletal muscle myoblasts which were first expanded under high mitogen conditions and later induced to differentiate by switching to a low serum medium (Trapnell et al set, **SI table S1**). A total of 96 cells were profiled with RNA-Seq at differentiation induction, as well as at 24h and 48h after medium switch, with a remaining 84 cells profiled at 72h. As expected, signaling entropy was highest in the myoblasts, with a stepwise reduction in signaling entropy observed at 24h (**Fig.4A**). No decrease in entropy was observed between 24 and 72h, indicating that commitment of cells to become differentiated skeletal muscle cells already happens early in the differentiation process. Over the whole timecourse, SCENT predicted a total of 3 potency states, with a distribution consistent with the time of sampling (**Fig.4B**). Cells sampled at differentiation induction were made up primarily of two potency states (**Fig.4C**, PS1 & PS2), which differed in terms of CDK1 expression, consistent with one subset (PS1) defining a highly proliferative subpopulation and with the rest (PS2) representing cells that have exited the cell-cycle (**SI fig.S8**). In total, SCENT predicted 4 landmarks, with one landmark defining undifferentiated (t=0) myoblasts of high potency (**Fig.4D**). Another landmark of lower potency contained cells at all time points, with cells expressing markers of mesenchymal cells (e.g PDFGRA and FN1/LTBP2) (**Fig.4D**). Cells from this landmark which were present at differentiation induction exhibited intermediate potency expressing low levels of CDK1 (**SI fig.S8 & Fig.4D**), suggesting that these are “contaminating” interstitial mesenchymal cells that were already present at the start of the time course, in line with previous observations [14, 15]. Importantly, SCENT correctly predicts that the potency of all these mesenchymal cells in this landmark does not change during the time-course, consistent with the fact that these cells are not primed to differentiate into skeletal muscle cells, but which nevertheless aid the differentiation process [14, 15]. Another landmark of intermediate potency predicted by SCENT defined a trajectory made up of cells expressing high levels of myogenic markers (*MYOG & IGF2*) from 24h onwards (**Fig.4D**). Thus, this landmark corresponds to cells that are effectively committed to becoming fully mature skeletal muscle cells. The final landmark consisted of cells exhibiting the lowest level of potency and emerged only at 48h, becoming most prominent at 72h (**Fig.4D**). As with the previous landmark, cells in this group also expressed myogenic markers, and likely represent a terminally differentiated and more mature state of skeletal muscle cells. In summary, SCENT inferred lineage trajectories that are highly consistent with known biology and with those obtained by previous algorithms such as Monocle [14] and MPath [15]. However, in contrast to Monocle and MPath, SCENT inferred these reconstructions without the explicit need of knowing the time-point at which samples were collected.

**Figure-4.**
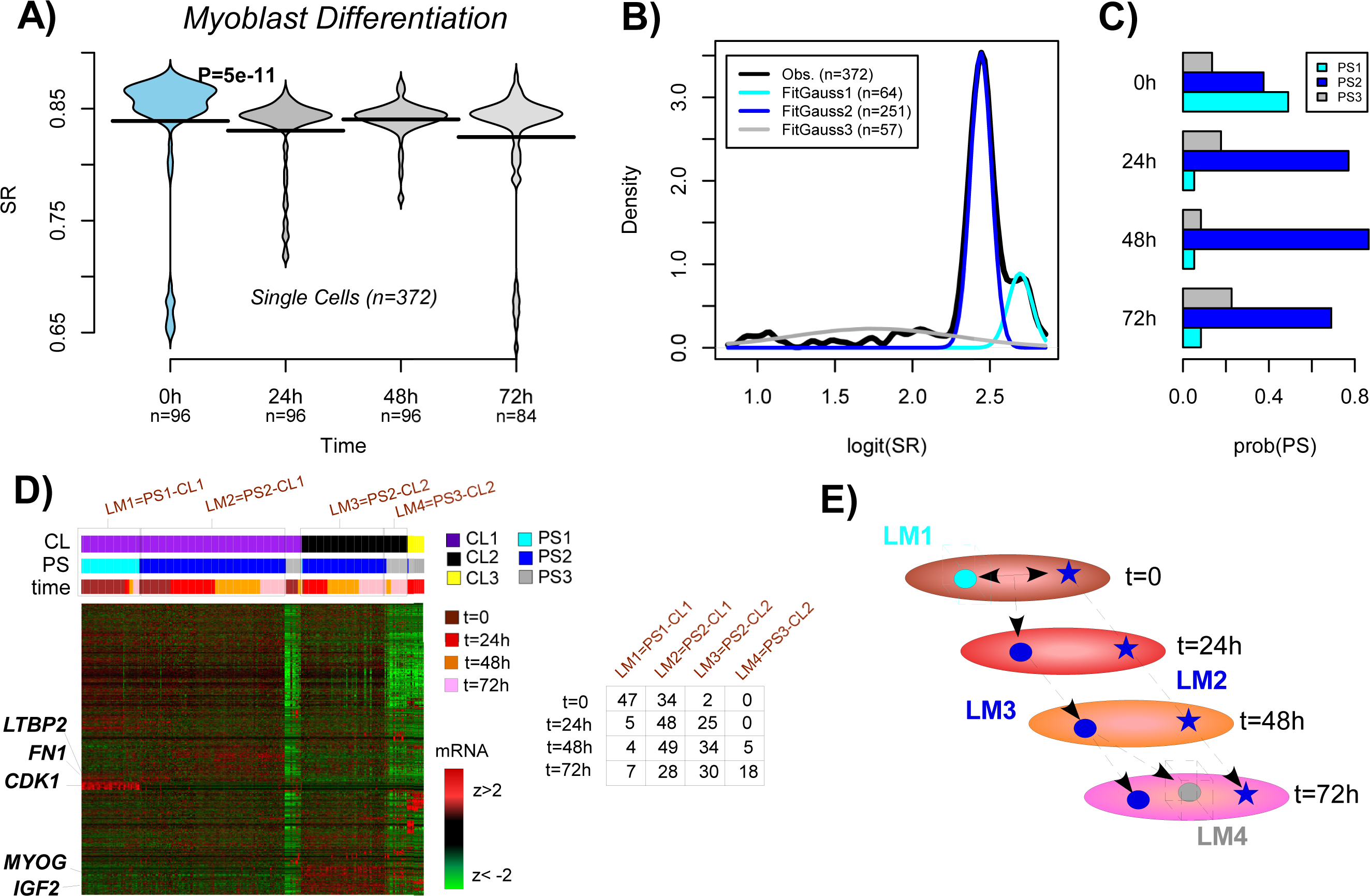
SCENT dissects distinct lineage trajectories in human myoblast differentiation. A) Signaling entropy (SR) vs. time point (0h, 24h, 48h, 72h) for a total of 372 single cells, collected during a time course differentiation experiment of human myoblasts (scRNA-Seq from Trapnell et al). Violin plots show the density distribution of SR values at each time point. P-value is from a one-tailed Wilcox rank sum test comparing timepoint 0h to 24h. **B)** SCENT Gaussian Model fit to SR values predicts 3 potency states (PS1, PS2, PS3). **C)** Probability distribution of potency states at each timepoint. **D)** Co-expression heatmap of highly variable genes obtained by SCENT predicting 3 main clusters. Single cells have been ordered, first by cluster, then by potency state and finally by their time of sampling, as indicated. Landmarks are indicated by rectangular boxes, and distribution of single cells across landmarks and timepoints is provided in table. Genes have been clustered using hierarchical clustering. Genes that are markers of the different landmarks have been highlighted. **E)** Inferred lineage trajectories between landmarks. Diagram illustrates an inferred two-phase trajectory, with one trajectory describing myoblasts of high potency (t=0, cyan circle) differentiating into skeletal muscle cells of intermediate potency (t=24 and 48) (blue circles) and a mixture of terminally differentiated and intermediate potency skeletal muscle cells (t=72) (grey and blue circle, respectively). A second trajectory/landmark describes a different cell-type (interstitial mesenchymal cells) whose intermediate potency state does not change during the time-course (blue stars).

### Signaling entropy detects drug resistant cancer stem cell phenotypes

Cancer cells are known to be less differentiated and to acquire a more plastic phenotype compared to non-malignant cells. Hence their signaling entropy should be higher than that of non-malignant cell-types. We confirmed this using scRNA-Seq data from 12 melanomas (Melanoma-set [22], **SI table S1**), for which sufficient normal and cancer cells had been profiled (**Fig.5A, SI fig.S9**). Although there was some variation in the signaling entropy of cancer cells between individuals, this variation was relatively small in comparison to the difference in entropy between cancer and normal cells. Combining data across all 12 patients, demonstrated a dramatic increase in the signaling entropy of single cancer cells compared to non-malignant ones (Wilcoxon rank sum test P<1e-500, **Fig.5B**).

**Figure-5.**
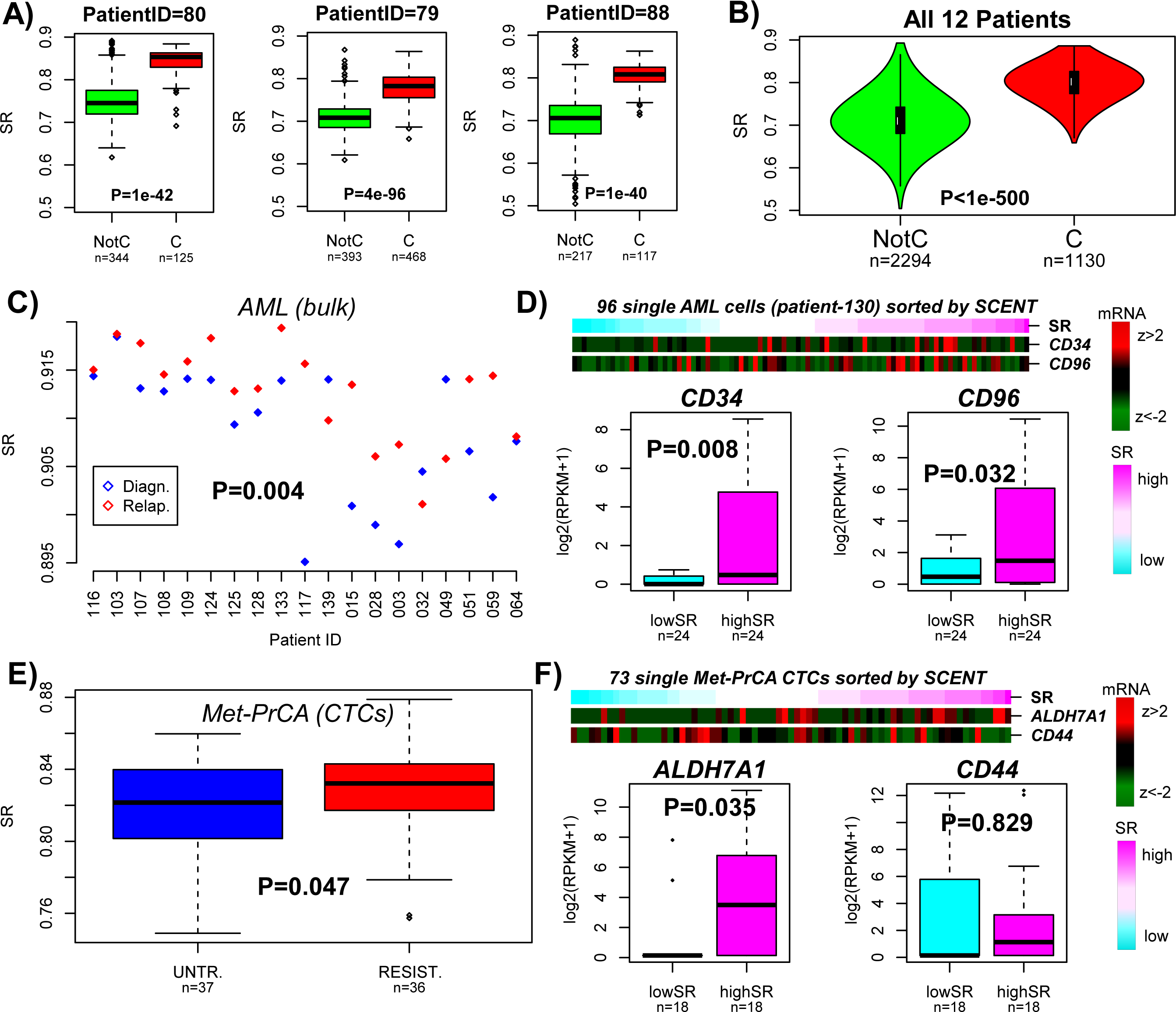
Increased signaling entropy in cancer cells and identification of drug resistant cancer stem cells. **A)** Boxplots of the signaling entropy (SR) for single melanoma cancer cells (C) compared to non-malignant (NotC) cells for 3 different melanoma patients (patient IDs given above each plot). Numbers of single cells are given below each boxplot. P-value is from a Wilcoxon rank sum test. **B)** As A), but now pooled across all 12 patients. **C)** Comparison of signaling entropy (SR) of 19 diagnostic acute myeloid leukemia bulk samples to relapsed samples from the same patients. Wilcox rank sum test P-value (one-tailed paired) is given. **D)** Sorting of 96 single AML cells from one patient according to signaling entropy and comparison of mRNA expression of AML CSC markers between low and high SR groups. P-values from a one-tailed Wilcox test. **E)** Comparison of signaling entropy (SR) of circulating tumor cells from metastatic prostate cancer patients who did not receive AR inhibitor treatment (UNTR) to those which developed resistance (RESIST). P-value from a one-tailed Wilcox test. **F)** Sorting of 73 single CTCs according to SCENT (signaling entropy, SR) into low and high SR groups. Correlation of gene expression of one putative CSC marker (ALDH7A1) with SR.

Since signaling entropy is increased in cancer and correlates with stemness, it could, in principle, be used to identify putative cancer stem cells (CSC) or drug resistant cells. To test this, we first computed and compared signaling entropy values for 38 acute myeloid leukemia (AML) bulk samples from 19 AML patients, consisting of 19 diagnostic/relapse pairs [31]. Confirming that signaling entropy marks drug resistant cell populations, we observed a higher entropy in the relapsed samples (paired Wilcox test P=0.004, **Fig.5C**). For one relapsed sample, scRNA-Seq for 96 single AML cells was available (AML set, **SI table S1**). We posited that comparing the signaling entropy values of these 96 cells would allow us to identify a CSC-like subset responsible for relapse. Since in AML there are well accepted CSC markers (CD34, CD96), we tested whether expression of these markers in high entropy AML single cells is higher than in low entropy AML single cells (**Fig.5D**). Both CD34 and CD96 were more highly expressed in the high entropy AML single cells (Wilcox test P=0.008 and 0.032, respectively, **Fig.5D**).

We next computed signaling entropies for 73 circulating tumor cells (CTCs) derived from 11 castration resistant prostate cancer patients (CTC-PrCa set, **SI table S1**), of which 5 patients exhibited progression under treatment with enzalutamide (an androgen receptor (AR) inhibitor) (n=36 CTCs), with the other 6 patients not having received treatment (n=37 CTCs) [32]. Although of marginal significance, signaling entropy was higher in the CTCs from patients exhibiting resistance (Wilcox test P=0.047, **Fig.5E**). Among putative prostate cancer stem cell markers (e.g. CD44, CD133, KLF4 and ALDH7A1) [32], we observed a positive association of signaling entropy with ALDH7A1 expression, suggesting that ADLH7A1 (and not other markers such as CD44) may mark specific prostate CSCs which are resistant to enzalutamide treatment (**Fig.5F**).

### Comparison of signaling entropy of bulk tissue to that of single cells reveals that intercellular expression heterogeneity is regulated

It has been proposed that expression heterogeneity of cell populations is regulated and optimized in a way which fulfills specific requirements such as pluripotency or homeostasis [3]. To test whether signaling entropy can predict such regulated expression heterogeneity, we compared the average of single-cell entropies with the signaling entropy of the bulk population. Specifically, we devised a “measure of regulated heterogeneity” (MRH), which measures the likelihood that the signaling entropy of the cell population could have been observed from picking a single cell at random from that population (**Online Methods, Fig.6A**). We first estimated MRH for the data from Chu et al, for which matched bulk and scRNA-Seq data was available. We first note that although for bulk samples entropy differences between cell-types were smaller, that they were nevertheless consistent with the trends seen at the single-cell level (**SI fig.S10 & Fig.2C**). The MRH for each of the six cell-types (hESCs, NPCs, DEPs, TBs, HFFs, ECs) in Chu et al, revealed evidence of regulated heterogeneity, with the entropy values of bulk samples being significantly higher than that of single-cells (**Fig.6B**). As a negative control, the signaling entropy of the average expression over bulk samples would not exhibit regulated heterogeneity since bulk samples are completely independent from each other (i.e. they are not linked in space or time and represent non-interacting cell populations). Confirming this, the MRH of the average expression taken over bulk samples, measured relative to individual bulk samples was not significant (Normal deviation test P=0.30, **Fig.6B**).

**Figure-6.**
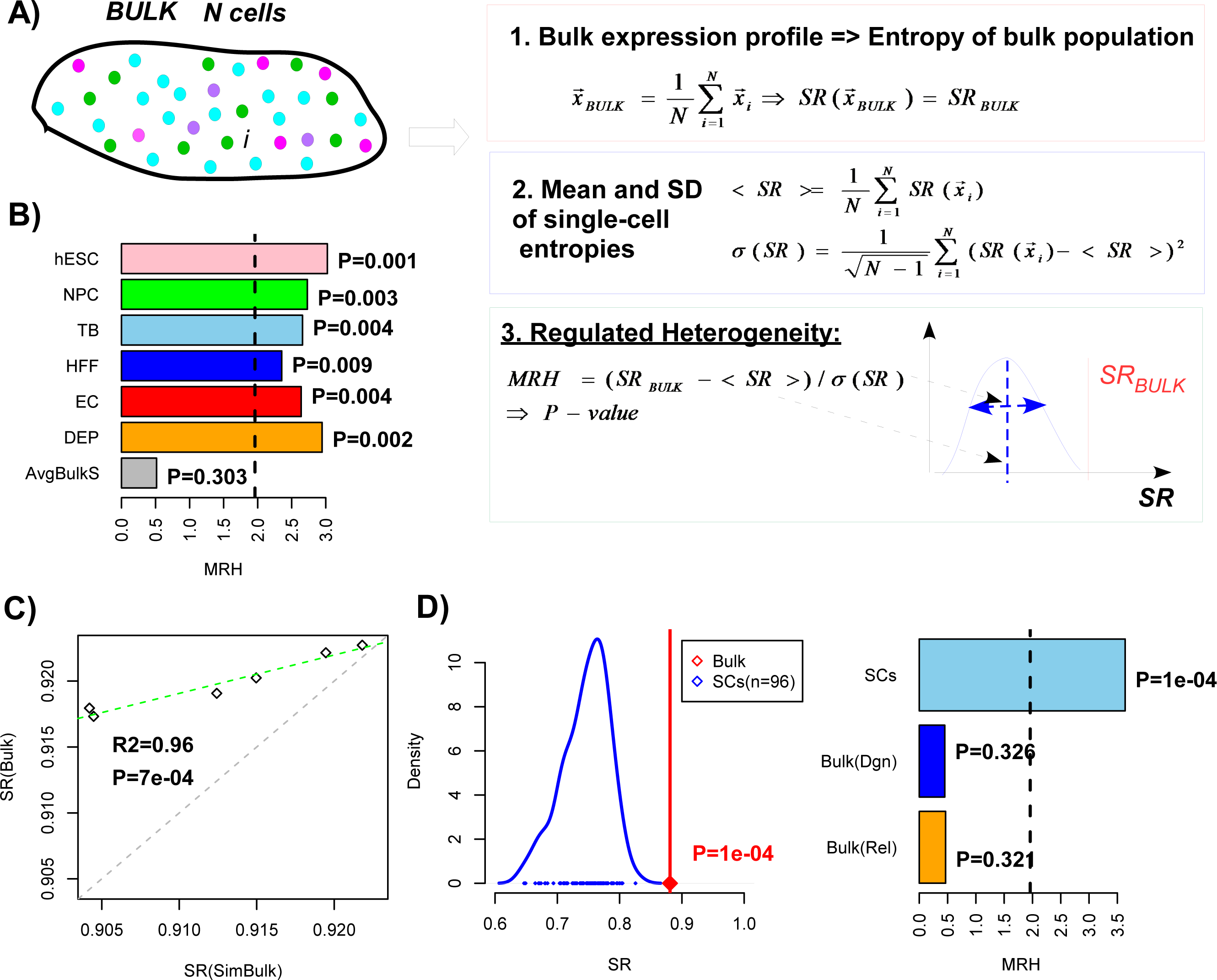
Signaling entropy predicts regulated expression heterogeneity of single-cell populations. **A)** Definition of the measure of regulated expression heterogeneity (MRH). The MRH is a z-statistic, obtained by measuring the deviation of the signaling entropy (SR) of the bulk expression profile from the mean of single-cell entropies, taking into account the variability of single-cell entropies in the population. **B)** Barplots of MRH for each cell-type population from Chu et al, representing the degree to which the signaling entropy of the cell population is higher than that of single-cells. P-values are from a one-tailed normal-deviation test. Dashed line indicates the line P=0.05. AvgBulkS compares the signaling entropy of the average expression over all bulk samples to that of the individual bulk samples, indicating that although the RHM is positive (signaling entropy increases), that it is not significantly higher than that of the individual bulk samples. **C)** Scatterplot of the signaling entropy of bulk samples (y-axis), representing 6 cell-types (hESCs, NPCs, DEPs, TBs, HFFs, ECs) against the corresponding signaling entropies of these cell populations obtained by first averaging the expression profiles of single-cells (“Simulated Bulk”, x-axis). R^2^ value and P-value are given with green dashed line representing the fitted regression. Observe how the signaling entropy of bulk samples is always higher than that obtained from first averaging expression of single cells, in line with the fact that the assayed single cells are a subpopulation of the full bulk sample. **D) Left panel:** Comparison of the signaling entropy of an acute myeloid leukemia (AML) bulk sample (red line and point) to the signaling entropies of 96 single AML cells (blue) from that bulk sample. P-value is from a one-tailed normal deviation test. **Right panel:** Comparison of the MRH value for the matched 96 single cells and bulk AML sample (SCs) to the MRH values obtained by comparing the signaling entropy of the average expression over 19 AML bulk samples to the signaling entropies of each individual AML bulk sample. The 19 AML bulk samples come in pairs, obtained at diagnosis (dgn) and relapse (rel), which are shown separately. P-values are from a one-tailed normality deviation test.

In order to obtain further evidence for regulated heterogeneity, we note that matched bulk RNA-Seq data is not absolutely required since bulk samples can be approximated by averaging the expression profiles of individual cells in the population. Indeed, we verified that the signaling entropy of the previous bulk samples correlated well with the entropy values obtained by averaging expression profiles of single cells, although as expected the values for the true bulk samples were always marginally higher, in line with the fact that the single cell assays only capture a subpopulation of the bulk sample (**Fig.6C**). Given this result, we explored if there is also regulated expression heterogeneity among normal cells of the tumor microenvironment using the average expression over single cells to approximate the bulk. This analysis was performed for T-cells and B-cells found in melanomas (Melanoma-set, **SI table S1**), for which sufficient numbers of single cells had been profiled. In all cases, signaling entropies of the bulk were much higher than expected based on the distribution of single-cell entropies (**SI fig.S11**). Evidence for regulated expression heterogeneity was also seen among the melanoma cancer cells from each of 12 patients (Combined Fisher test P<1e-6, **SI fig.S12**). We also analysed RNA-Seq data for 96 single cancer cells from a relapsed patient with acute myeloid leukemia (AML) (AML set [31], **SI table S1**). The signaling entropy for the AML cell population was 0.88, significantly larger than the maximal value over the 96 cells (SR=0.82, Normal deviation test P<0.001, **Fig.6D**). Again, to illustrate that this regulated heterogeneity is a result of inter-cellular interactions at the single-cell level, we analysed all 19 bulk AML samples at relapse, treating bulk samples from independent AML patients as if they were single cells from a common population. Estimating the signaling entropy of the average expression profile over all 19 bulk samples did not reveal a value significantly higher than that of the individual bulk samples (Normal deviation test P=0.32, **Fig.6D**). This result was unchanged if the bulk samples at relapse were replaced with bulk samples at diagnosis (**Fig.6D**). In summary, these data strongly support the view that the differentiation potential or phenotypic plasticity of a cell population is higher than that of a randomly picked single cell in the population, consistent with a model in which expression heterogeneity between single cells is regulated.

## Discussion

Although Waddington proposed his famous epigenetic landscape of cellular differentiation many decades ago [10], it has proved challenging to construct a robust molecular correlate of a cell’s elevation in this landscape. Here we have made significant progress, demonstrating that the differentiation potency and phenotypic plasticity of single cells, be they normal or malignant, can be estimated *in-silico* from their RNA-Seq profile using signaling entropy. As we have seen, signaling entropy can accurately discriminate pluripotent from multipotent and differentiated cells, without the need for feature selection or training, outperforming a pluripotency gene expression signature and providing a *more general* measure of differentiation potency.

The ability of signaling entropy to independently order single cells according to differentiation potency is a central component of the SCENT algorithm, which, as shown here, can help quantify and identify biologically relevant intercellular expression heterogeneity and cell subpopulations. Indeed, key findings which strongly support the validity of SCENT are the following: (i) using SCENT we were able to correctly predict that a hESC population contains a small fraction of cells of lower potency which are primed for differentiation, (ii) SCENT inferred that an assayed neural progenitor cell population was made up two distinct subsets, correctly predicting that only the lower potency subset represents bona-fide NPCs (as determined by expression of known neural stem cell markers), (iii) in a time course differentiation experiment of human myoblasts, SCENT correctly identified a contaminating interstitial mesenchymal cell population, *whose potency did not change appreciably during the experiment*. The ability of SCENT to assign single cells and cell subpopulations to specific potency states thus adds novel insight and functionality over what can be achieved with other algorithms such as Monocle or MPath. Alternatively, signaling entropy could be combined with existing algorithms like Monocle to empower their inference, since signaling entropy provides an unbiased, independent, approach to ordering of single cells in pseudo-time, i.e. it constitutes an approach which does not need to know the time point or nature of the assayed cells.

In a proof of principle analysis, we further demonstrated the ability of SCENT to identify putative drug resistant cancer stem cells, encompassing two different cancer-types (AML and prostate cancer), including CTCs. The ability to quantify stemness in cancer cell populations, either in tissue or in circulation, is a task of enormous importance. As shown here, as well as in our previous work on bulk cancer tissue [9, 11, 13], signaling entropy is, so far, the only single sample measure to have been conclusively demonstrated to robustly correlate with both stemness and cancer. Indeed, a recent study by Gruen et al [18] explored a very different measure of transcriptome entropy, but which was not demonstrated to correlate well with differentiation potency or cancer. Likewise, signaling entropy is a more general measure of stemness/plasticity outperforming existing pluripotency expression signatures, as shown here and previously [11].

Importantly, signaling entropy also provides a computational framework in which to understand differentiation potency at the macroscopic (cell population) level from the corresponding potencies of single cells. As shown here, signaling entropy of cell populations, be they normal or malignant cells, exhibit synergy, with the entropy of the bulk being substantially higher than the entropy values of single cells. While no existing assay can measure all single cells in a population, we nevertheless demonstrated that our result is non-trivial, since mixing up bulk samples (to serve as a negative control) did not reveal such synergy. Biologically, increased potency of a cell population as a result of synergistic cell-cell interactions, supports the view that features such as pluripotency are best understood at the cellular population level [3].

To conclude, signaling entropy and the SCENT algorithm provide a computational framework to advance our understanding of single-cell biology. We envisage that SCENT will be of great value for quantifying biologically relevant intercellular heterogeneity and identifying key cell subpopulations in scRNA-Seq experiments.

## Online Methods

### Single cell and bulk RNA-Seq data sets

The main datasets analysed here, the NGS platform used and their public accession numbers are listed in **SI table-1**. Below is a more detailed description of the samples in each data set:

*Chu et al Set:* This RNA-Seq dataset derives from Chu et al [27]. This set consisted of 4 experiments. Experiment-1 generated scRNA-Seq data for 1018 single cells, composed of 374 hESCs (212 single-cells from H1 and 162 from H9 cell line), 173 neural progenitor cells (NPCs), 138 definite endoderm progenitors (DEPs), 105 mesoderm derived endothelial cells (ECs), 69 trophoblast cells (TBs), 159 human foreskin fibroblasts (HFFs). Experiment-2 is a time-course differentiation of single-cells, specifically of hESCs induced to differentiate into the definite endoderm, via a mesoendoderm intermediate. Timepoints assayed were before induction (t=0h, n=92), 12 hours after induction (12h, n=102), 24h (n=66), 36h (n=172), 72h (n=138) and 96h (n=188). Experiment-3 matches experiment-1 and consists of RNA-Seq data from 19 bulk samples: 7 representing hESCs, 2 representing NPCs, 2 TBs, 3 HFFs, 3 ECs and 2 DEPs. Experiment-4 consists of 15 RNA-Seq profiles from bulk samples, profiled as part of the time-course differentiation experiment (Experiment-2), with 3 samples per time-point (12h, 24h, 36h, 72h, 96h).

*Melanoma Set:* This scRNA-Seq dataset derives from Tirosh et al [22], and consists of 4645 single-cells derived from the tumor microenvironment of 19 melanoma patients. Of these, 3256 are non-malignant cells, encompassing T-cells (n=2068), B-cells (n=515), Natural Killer cells (n=52), Macrophages (n=126), Endothelial Cells (EndC, n=65) and cancer-associated fibroblasts (CAFs, n=61). The rest of single cells profiled were malignant melanoma cells (n=1257).

*AML Set:* This set derives from Li et al [31]. A total of 96 single cells from a relapsed acute myeloid leukemia (AML) patient (patient ID=130) were profiled. In addition, 38 paired bulk AML samples were profiled from 19 patients (all experiencing relapse), with 19 samples obtained at diagnosis and with the other matched 19 samples obtained at relapse.

*hESC Set:* This set derives from Yan et al [28]. It consists of 124 single cell profiles, of which 90 are from different stages of embryonic development, with 34 cells representing hESCs. These 34 hESCs were derived from the inner cell mass, with 8 cells profiled at primary outgrowth and 26 profiled at passage-10. The 90 single cells from the pre-implantation embryo were distributed as follows: Oocyte (n=3), Zygote (n=3), 2-cell embryo (n=6), 4-cell embryo (n=12), 8-cell embryo (n=20), morulae (n=16), late blastocyst (n=30).

*Trapnell et al set:* This scRNA-Seq set derives from Trapnell et al [14]. It consists of a timecourse differentiation experiment of human myoblasts, which profiled a total of 372 single cells: 96 cells at t=0 (time at which differentiation was induced), 96 at t=24h after induction, another 96 at t=48h after induction, and 84 cells at 72h post-induction.

*CTC-PrCa set:* This scRNA-Seq dataset derives from Miyamoto et al [32].We focused on a subset of 73 single-cells from castration resistant prostate cancers, of which 36 derived from patients who developed resistance to enzulatamide treatment, with the remaining 37 derived from treatment-naïve patients.

### The Single-Cell Entropy (SCENT) algorithm

There are five steps to the SCENT algorithm: (1) Estimation of the differentiation potency of single cells via computation of signaling entropy, (2) Inference of the potency state distribution across the single cell population, (3) Quantification of the intercellular heterogeneity of potency states, (4) Inference of single cell landmarks, representing the major potency-coexpression clusters of single cells, (5) Lineage trajectory (or dependency network) reconstruction between landmarks. We now describe each of these steps:

#### 1. Computation of signaling entropy

The computation of signaling entropy for a given sample proceeds using the same prescription as used in our previous publications [9, 11]. Briefly, the normalized genome-wide gene expression profile of a sample (this can be a single cell or a bulk sample) is used to assign weights to the edges of a highly curated protein-protein interaction (PPI) network. The construction of the PPI network itself is described in detail elsewhere [11], and is obtained by integrating various interaction databases which form part of Pathway Commons (www.pathwaycommons.org) [33]. The weighting of the network via the transcriptomic profile of the sample provides the biological context. The weight of an edge between protein *g* and protein *h,* denoted by *w*_*gh*_, is assumed to be proportional to the normalized expression levels of the coding genes in the sample, i.e. we assume that *w*_*gh*_ ~ *x*_*g*_ *x*_*h*_. We interpret these weights (if normalized) as interaction probabilities. The above construction of the weights is based on the assumption that in a sample with high expression of *g* and *h,* that the two proteins are more likely to interact than in a sample with low expression of *g* and/or *h.* Viewing the edges generally as signaling interactions, we can thus define a random walk on the network, assuming we normalize the weights so that the sum of outgoing weights of a given node *i* is 1. This results in a stochastic matrix, *P*, over the network, with entries

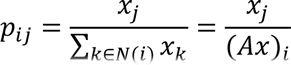

where *N(i)* denotes the neighbors of protein *i*, and where *A* is the adjacency matrix of the PPI network (*A*_*ij*_=1 if *i* and *j* are connected, 0 otherwise, and with *A*_*ii*_=0). The signaling entropy is then defined as the entropy rate (denoted *Sr*) over the weighted network, i.e.

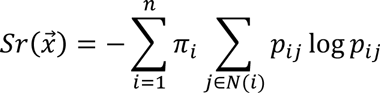

where *π* is the invariant measure, satisfying *πP=π* and the normalization constraint *π*^*T*^**1**=1.

Assuming detailed balance, it can be shown that π_i_ = *x*^*i*^(Ax)^*i*^/(*x*_*T*_*Ax*). Given a fixed adjacency matrix *A* (i.e. fixing the topology), it can be shown that the maximum possible *Sr* among all compatible stochastic matrices *P*, is the one with 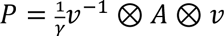 where 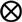 denotes product of matrix entries and where *v* is the dominant eigenvector of *A*, i.e. *Av=λv* with *λ* the largest eigenvalue of *A*. We denote this maximum entropy rate by *maxSr*, and define the normalized entropy rate (with range of values between 0 and 1) as

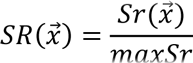

Throughout this work, we always display this normalized entropy rate.

As shown by us previously, signaling entropy is influenced mainly by the invariant measure *π*, since the dynamic range of local signaling entropies S^*i*^ = − ∑_*j∈N(i)*_ *p*_*ij*_ log *p*_*ij*_ is in practice quite small [12]. In a mean field approximation, it is clear that 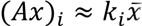, where 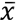 is the average expression over all genes in the network. Thus, 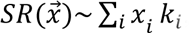, i.e, the signaling entropy is effectively the Pearson correlation of the cell’s transcriptome and the connectome from the PPI network. In this work, although we never use this approximation, in practice this approximation is highly accurate and helps understand the biological features of signaling entropy [12].

#### 2. Inference of potency states

In this work, we show that signaling entropy (i.e. the entropy rate *SR*) provides a proxy to the differentiation potential of single cells. We can model a cell population as a statistical mechanical model, in which each single cell has access to a number of different potency states. For a large collection of single cells we can estimate their signaling entropies, and infer from this distribution of signaling entropies the number of underlying potency states using a mixture modeling framework. Since *SR* is bounded between 0 and 1, we first conveniently transform the *SR* value of each single cell into their logit-scale, i.e. *y(SR)=log_2_(SR/(1-SR))*. Subsequently, we fit a mixture of Gaussians to the *y(SR)* values of the whole cell population, and use the Bayesian Information Criterion (BIC) (as implemented in the *mclust* R-package) [34] to estimate the optimal number *K* of potency states, as well as the state-membership probabilities of each individual cell. Thus, for each single cell, this results in its assignment to a specific potency state.

#### 3. Quantifying intercellular heterogeneity of potency states

For a population of *N* cells, we can then define a probability distribution *pk* over the inferred potency states. For *K* inferred potency states, one can then define a normalized Shannon Index (*SI*):

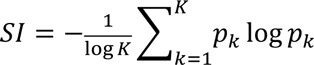

which measures the amount of heterogeneity in potency within the single-cell population (1=high heterogeneity in potency, 0=no heterogeneity in potency).

#### 4. Inference of co-expression clusters and landmarks

With each cell assigned to a potency state, we next perform clustering (using the scRNA-seq profiles) of the single cells. We use the Partitioning-Around-Medoids (PAM) algorithm with the average silhouette width to estimate the optimal number of clusters, a combination which was found to be among the most optimal clustering algorithms in applications to omic data [35]. Clustering of the cells is performed over a filtered set of genes that are identified as those driving most variation in the complete dataset, as assessed using SVD. In detail, we perform a SVD on the full z-scored normalized RNA-seq profiles of the cells, selecting the significant components using RMT [36] and picking the top 5% genes with largest absolute weights in each significant component. The final set of genes is obtained by the union of those identified from each significant componente. PAM-clustering (with a Pearson distance correlation metric) of all cells results in the assignment of each cell into a co-expression cluster, with a total number of *n*_*p*_ cell-clusters. Thus, each cell is assigned to a unique potency state and co-expression cluster. Finally, landmarks are identified by selecting potency-state cluster combinations containing at least 1 to 5% of all single cells. Importantly, each of these landmarks has a specific potency state and mean signaling entropy value, allowing ordering of these landmarks according to potency.

#### 5. Inference of lineage trajectories

For each landmark in step-4, we compute centroids of gene expression using only cells that are contained within that landmark and defined only over the genes used in the PAM-clustering. Partial correlations [37, 38] between the centroid landmarks are then estimated to infer trajectories/dependencies between landmarks. Significant positive partial correlations may indicate transitions between landmarks. Since each landmark has a signaling entropy value associated with it, directionality is inferred by comparing their respective potency states.

**Software Availability:** SCENT is freely available as an R-package from github:

https://github.com/aet21/SCENT

### Estimation of cell-cycle and TPSC pluripotency scores

To identify single cells in either the G1-S or G2-M phases of the cell-cycle we followed the procedure described in [22]. Briefly, genes whose expression is reflective of G1-S or G2-M phase were obtained from [29, 30]. A given normalized scRNA-Seq data matrix is then z-score normalized for all genes present in these signatures. Finally, a cycling score for each phase and each cell is obtained as the average z-scores over all genes present in each signature.

To obtain an independent estimate of pluripotency we used the pluripotency gene expression signature of Palmer et al [21], which we have used extensively before [11]. This signature consists of 118 genes that are overexpressed and 39 genes that are underexpressed in pluripotent cells. The TPSC score for each cell with scRNA-Seq data is obtained as the t-statistic of the gene expression levels between the overexpressed and underexpressed gene categories. Optionally, the scRNA-Seq is z-score normalized beforehand and the t-statistic is obtained by comparing expression z-scores. However, we note that the z-score procedure uses information from all single cells, so the fairest comparison to signaling entropy means we ought to compare expression levels. We note that the TPSC scores obtained from z-scores or expression levels were highly correlated and did not affect any of the conclusions in this manuscript.

## Supplementary Material

All Supplementary Tables and Figures can be found in the Supplementary Information document.

## Author Contributions

Manuscript was conceived and written by AET. Statistical analyses were performed by AET.

## Acknowledgements

This work was supported by NSFC (National Science Foundation of China) grants, grant numbers 31571359 and 31401120 and by a Royal Society Newton Advanced Fellowship (NAF project number: 522438, NAF award number: 164914). The author also wishes to thank Tariq Enver and Guo-Cheng Yuan for stimulating discussions.

